# Elucidating gene expression patterns across multiple biological contexts through a large-scale investigation of transcriptomic datasets

**DOI:** 10.1101/2022.01.18.476735

**Authors:** Rebeca Queiroz Figueiredo, Sara Díaz del Ser, Tamara Raschka, Martin Hofmann-Apitius, Alpha Tom Kodamullil, Sarah Mubeen, Daniel Domingo-Fernández

## Abstract

Distinct gene expression patterns within cells are foundational for the diversity of functions and unique characteristics observed in specific contexts, such as human tissues and cell types. Though some biological processes commonly occur across contexts, by harnessing the vast amounts of available gene expression data, we can decipher the processes that are unique to a specific context. Therefore, with the goal of developing a portrait of context-specific patterns to better elucidate how they govern distinct biological processes, this work presents a large-scale exploration of transcriptomic signatures across three different contexts (i.e., tissues, cell types, and cell lines) by leveraging over 600 gene expression datasets categorized into 98 subcontexts. The strongest pairwise correlations between genes from these subcontexts are used for the construction of co-expression networks. Using a network-based approach, we then pinpoint patterns that are unique and common across these subcontexts. First, we focused on patterns at the level of individual nodes and evaluated their functional roles using a human protein-protein interactome as a referential network. Next, within each context, we systematically overlaid the co-expression networks to identify specific and shared correlations as well as relations already described in scientific literature. Additionally, in a pathway-level analysis, we overlaid node and edge sets from co-expression networks against pathway knowledge to identify biological processes that are related to specific subcontexts or groups of them. Finally, we have released our data and scripts at https://zenodo.org/record/5831786 and https://github.com/ContNeXt/, respectively and developed ContNeXt (https://contnext.scai.fraunhofer.de/), a web application to explore the networks generated in this work.

## 1. Introduction

While gene expression profiling has markedly improved our understanding of the molecular underpinnings of biological processes, the knowledge we acquire from a particular study performed within a given context may not generalize to another. For instance, accumulating evidence shows that average gene expression varies extensively across cell lines or tissues of the same organism (Sonawane *et al*., 2017; Whitehead and Crawford, 2005) as well as across species (Romero *et al*., 2012). Context-specificity has also been noted when investigating the reproducibility of protein-protein interactions (PPIs) across conditions in literature-curated PPI databases in Stacey *et al*. (2018), finding no evidence for the occurrence of anywhere from 19 to 55% of interactions reported in these databases. These findings, however, are not altogether surprising given that PPI databases often store interactions that occur across various experimental conditions and contexts which may fail to be observed if either of these were to vary. Crucially, it is often these context-specific differences which are responsible for the variability of functions and unique characteristics of diverse cell types and tissues and their investigation is thus fundamental in understanding human biology.

Gene expression patterns that are specific to certain cell types or tissues can help us to better understand normal human physiology (e.g., which biological processes occur in specific cell types or tissues) as well as development biology (e.g., which genes are expressed in specific cell types or tissues at various developmental stages), and several studies have investigated differences in these two contexts. Specifically, Pierson *et al*. (2015) and Dobrin *et al*. (2009) analyzed gene expression patterns at the tissue-level, revealing function-specific patterns and subnetworks associated with obesity. Similarly, McKenzie *et al*. (2018) analyzed co-expression changes in different cell types of the brain, discovering significant cell type-specific expression signatures, while also finding well-known cell type marker genes among the most enriched genes across cell types.

Another relevant context is cell line information, as these are widely used for the study of biological processes. In particular, cancer cell lines, such as HeLa, are frequently employed, having had many interactions characterized on them and representing the foremost models for the study of cancer biology as well as numerous other disease and normal conditions. Nonetheless, even cell lines classified to the same tissue can exhibit significant differences in gene expression (Lee *et al*. 2018). For example, a study by Yu *et al*. (2019) found that certain cell lines may not resemble the primary cells from which they originated. The discrepancies in regulation patterns across specific cell lines deem it necessary to employ tools such as the CellExpress system (developed by Lee *et al*. (2018) which enables the analysis of over 4,000 cancer cell lines for differences in gene expression levels) and resources such as the TCGA-110-CL cell line panel (Yu *et al*. 2019) to identify which cell lines are more suitable for a given study.

Biological networks of different types can be used to represent patterns characteristic to a particular context. These context-specific networks can be categorized based on whether they are directly derived from knowledge or data. Rachlin *et al*. (2005) and Stacey *et al*. (2018) are two illustrations of knowledge-driven approaches where authors generated context-specific PPI networks by respectively leveraging information about biological processes from GO (The Gene Ontology Consortium *et al*., 2021) and co-occurrence literature. Similarly, the analysis of transcriptomic data through the construction of gene co-expression networks (Langfelder *et al*., 2008) can also serve to better understand context-specific patterns within datasets (Oldham *et al*., 2008; Farahbod and Pavlidis, 2020). Finally, hybrid approaches, as demonstrated by Kitsak *et al*. (2016), have leveraged gene expression data from 64 different tissues and mapped genes expressed in specific tissues to a protein-protein interactome, revealing that these context-specific genes tend to be located in close proximity within the interactome. Furthermore, in Sonawane *et al*. (2017), the authors describe a method to combine gene co-expression data with PPIs as well as interactions between transcription factors (TFs) and gene targets to identify tissue-specific network nodes and edges. Their study reveals that TF expression is less indicative of tissue specificity than the genes which encode for them and suggests this specificity arises from altered TF gene targeting rather than increased targeting. It is important to note that while transcriptomic experiments are often used as a proxy to reflect protein expression, the correlation between the two is often below 0.5 on average (Nusinow *et al*., 2020; Trapotsi *et al*., 2022). Nevertheless, correlations between differentially expressed genes (DEGs) and their protein products have been shown to be significantly higher than in non-DEGs, which suggests that stronger transcriptomic signals are reflected in protein translation (Koussounadis *et al*., 2015).

One of the challenges in conducting these hybrid approaches (i.e., approaches that combine data- and knowledge-derived networks) is the limited availability of context-specific resources on a large-scale (e.g., hundreds of experiments conducted within the same or similar conditions or context-specific interactomes). While there are several co-expression databases dedicated to storing context-specific information, such as species (Obayashi *et al*., 2019 and Lee *et al*., 2020), the vast majority of transcriptomic datasets are not annotated with context information and thus, cannot be systematically leveraged to conduct contextualized analyses on a large-scale. Nonetheless, the Gemma system (Lim *et al*., 2021) has been made available to provide thousands of curated datasets; thus, more easily enabling data reuse and secondary analyses.

In this work, we apply a network-based approach to investigate transcriptomic patterns observed in a variety of subcontexts classified under three major biological contexts (i.e., tissues, cell types, and cell lines) by leveraging over 600 gene expression datasets **(Figure 1A)**. To do so, we first construct co-expression networks that capture the strongest gene expression correlations observed in each subcontext **(Figure 1B)**. Subsequently, a series of network-based analyses are conducted to enable the exploration of the similarities and differences across co-expression networks and provide insights on gene co-expression patterns across contexts **(Figure 1C)**. Furthermore, we study the consensus between patterns identified in the co-expression network and a human protein-protein interactome as well as pathways knowledge. Finally, we present ContNeXt, a web application we have developed to enable researchers to explore and reuse our work.

**Figure 1.**
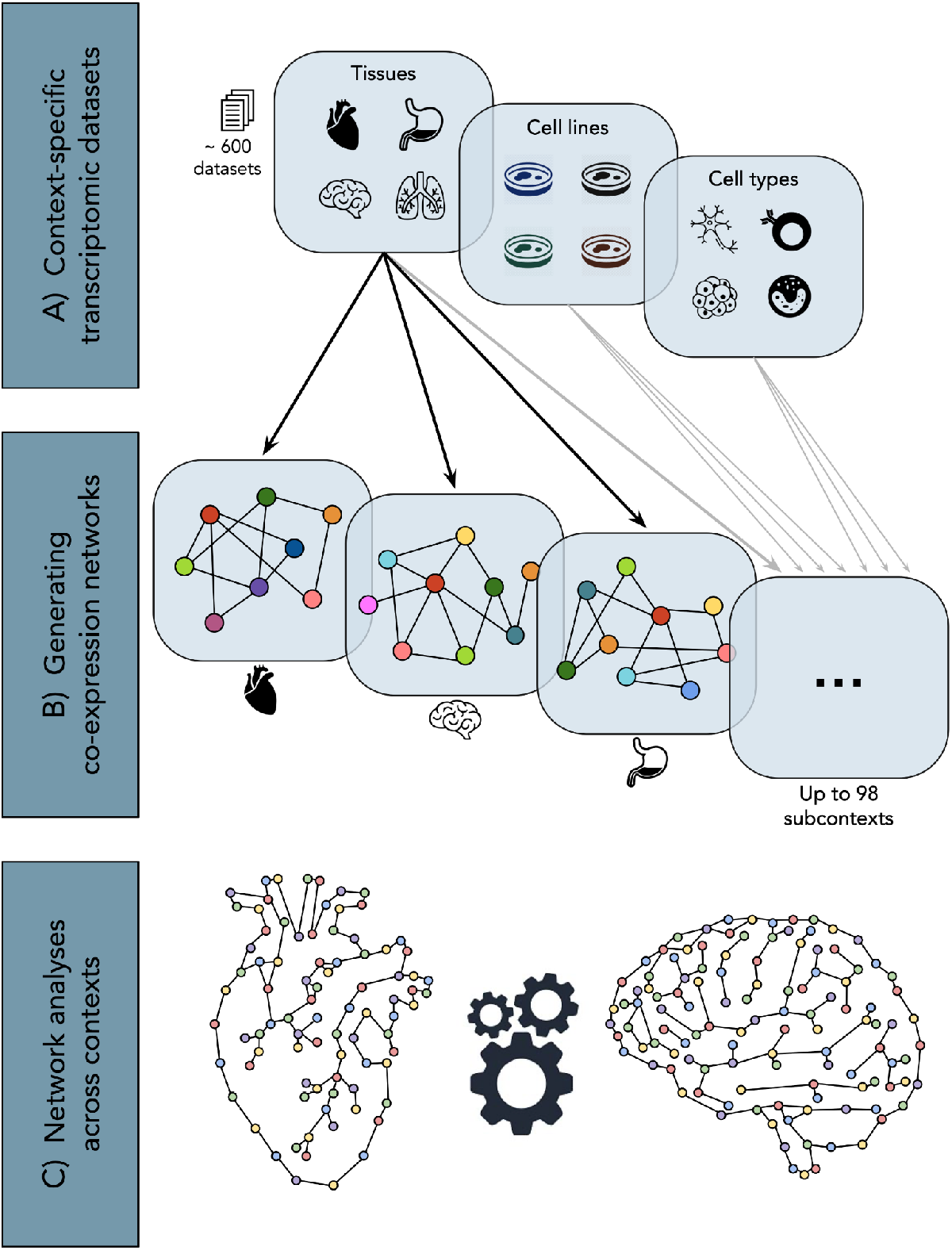
Conceptualization of the presented study. **A)** Over 600 context-specific transcriptomic datasets are collected and classified into 98 subcontexts (e.g., heart, astrocyte, and HeLa cell) under 3 major contexts (i.e., tissues, cell types, and cell lines), leveraging the Gemma database (Zoubarev *et al*., 2012; Lim *et al*., 2021) **B)** Co-expression networks comprising the most strongly correlated edges observed in each subcontext are generated. **C)** Network analyses provide insights on both common and unique patterns across the multiple contexts studied.

## 2. Methodology

### 2.1. Gene expression datasets

We identified publicly available transcriptomic datasets from each of the three contexts evaluated (i.e., tissues, cell types, and cell lines) using Gemma, a manually curated database containing metadata for over 10,000 datasets (Zoubarev *et al*., 2012; Lim *et al*., 2021) **(Figure 1A)**. This metadata is programmatically accessible through Gemma’s API (https://gemma.msl.ubc.ca/resources/restapidocs) and is annotated using different ontologies. Specifically, for each of the three contexts of interest, the following ontologies were used: i) UBERON for tissues (Mungall *et al*., 2012), ii) Cell Ontology (CL) for cell types (Diehl *et al*., 2016), and iii) Cell Line Ontology (CLO) for cell lines (Sarntivijai *et al*., 2014).

Leveraging the metadata from Gemma, we were able to classify the samples from each dataset to their corresponding context(s). To guarantee the quality of the annotations, we conducted an additional manual curation step where we confirmed that the Gemma sample annotations matched an ontology term for the given context present in the metadata, if available. Additionally, we filtered out samples that were not control or reference samples as our work focuses on comparing a normal physiological state in a variety of contexts. Finally, Gemma also includes annotations on dataset quality and samples that were annotated as unusable were excluded from our study.

After the initial annotation and curation steps, we implemented scripts for the downloading and processing of datasets found in Gene Expression Omnibus (GEO) (Edgar *et al*., 2002). While GEO incorporates several platforms, each measures different transcripts and requires a dedicated pipeline, and merging data from several platforms is a complicated task which can introduce biases from probe sequences, arrays, or laboratory effects. Furthermore, conducting analyses combining raw data from multiple platforms can also introduce biases (Rung and Brazma, 2013). Thus, our work focuses on the most commonly used platform for humans, the Affymetrix GeneChip Human Genome U133 Plus 2.0 Array platform (accession on GEO: GPL570). Out of 10,388 datasets in Gemma as of 22/04/2021, 9,778 were filtered out while 610 remained for any one of the three contexts. In total, the tissue context was divided into 46 subcontexts, while the cell line and cell type contexts each contained 22 and 30 subcontexts, respectively **(see Supplementary Tables 1-3)**.

### 2.2. Generating co-expression networks from gene expression data

Co-expression networks were constructed using the WGCNA package in R (Langfelder *et al*., 2008). We followed the same procedure outlined in our previous work (Figueiredo *et al*., 2021) to define the co-expression networks **(Figure 1B)**. This procedure focuses on the 1% highest similarity in the topological overlap matrix (TOM) to define the co-expression network for each subcontext; thus, facilitating the comparison of networks of the same size using a conservative cut-off in benchmark studies (Perkins and Langston, 2009). Given the platform used in this study, the most similar 1% in the TOM corresponds to 2,036,667 edges. We would like to note that the 1% cut-off is required as otherwise the networks would be fully connected, while we intend to focus only on the edges representing the most relevant transcriptomic patterns observed within each context. As edges representing a high topological overlap are also highly correlated in the TOM, we interchangeably refer to these edges as correlations for simplicity. Although this is not precise, the TOM value is based on the signed correlation but also takes the connectedness of nodes into account.

To run WGCNA, we used the raw expression data in the form of .CEL-files. Each dataset was individually pre-processed with the RMA function of the *oligo* R package to conduct background subtraction and quantile normalization. Next, we merged all samples from different datasets that belong to the same subcontext and applied batch correction using ComBat (Johnson *et al*., 2007). Regarding the mapping of the probes to genes, if there were multiple probes mapping to the same gene, we kept the most variable probe.

### 2.3. Protein – protein interaction network

We built a human protein-protein interactome as described in our previous work (Figueiredo *et al*., 2021) as a knowledge template to compare against the co-expression networks generated. The interactome comprises interactions from well-established databases, including KEGG (Kanehisa *et al*., 2021) and Reactome (Jassal *et al*., 2020). This network aims at representing the set of interactions that can occur in a physiological context, though it is worth mentioning that each of these interactions may not necessarily be occurring in a particular context at any given time.

### 2.4. Analyses

#### 2.4.1. Controllability analysis

One of the more advanced techniques in analyzing networks is examining its controllability. We employed an algorithm developed by Liu *et al*. (2011) which explores control theory to study the controllability of a directed network and thus identify driver nodes (i.e., the set of nodes that can offer control over the whole network) in order to classify each node and edge in a network as indispensable, dispensable, or neutral. Ideally, minimizing the number of driver nodes offers adequate control over the network regarding the given biological system’s dynamics. Using this algorithm, both nodes and edges can be classified as indispensable, dispensable, or neutral if their removal creates the need to increase, decrease, or cause no change in the number of driver nodes, respectively, so that controllability is maintained.

#### 2.4.2. Pairwise co-expression network similarity

To evaluate similarity across co-expression networks, we calculated the overlap of edges across each pair of co-expression networks within a given context. Since all co-expression networks have the same number of edges, the number of shared edges between networks is readily comparable without the need to normalize values.

#### 2.4.3. Similarity between co-expression networks and the interactome

We assessed the similarity of each co-expression network to the human interactome by calculating the number of shared edges. Here, it is important to note that edge directionality is ignored in the interactome since co-expression networks are inherently undirected. Furthermore, we evaluated the significance of the overlap by comparing the interactome to 1,000 permuted co-expression networks. Permuted versions of the co-expression networks were created using the XSwap algorithm (Hanhijärvi *et al*., 2009) (source code available at https://github.com/hetio/xswap), which ensures that the permuted versions preserve the structure of the original network (i.e., all edges are shuffled while maintaining the degree of each node).

#### 2.4.4. Pathway – co-expression network similarity

To investigate the correspondence of transcriptomic signatures from co-expression networks with pathway knowledge, each of the context-specific co-expression networks were overlaid with pathways from KEGG (Kanehisa *et al*., 2021). The KEGG database was exclusively employed as it contains a feasible number of pathways for analysis (i.e., less than 350). For each gene set of a given pathway *P* from KEGG, we calculate every pairwise combination of nodes (*C*_*n*_ ) in *P* to determine the fraction of node combination pairs in *C*_*n*_ that exist as an edge in a given co-expression network *N* = (*n*’, *E*_*N*_) where *n’* is the set of nodes in the co-expression network and *E*_*N*_ is the set of edges which connect the nodes *n’*. We term this the edge overlap, where *edge overlap* = |{ ∀*e*_*u,v*_ *s. t*. (*u, v*) ∈ *C*_*n*_ Λ *u, v* ∈ *n*’ Λ *e*_*u,v*_ ∈ *E*_*N*_}|. The proportion of *C*_*n*_ that is in the edge overlap is the pathway-network similarity **(Equation 1)**. Using the pathway-network similarity, we create a similarity matrix with each network of a given context against every pathway from KEGG. This matrix is subsequently used to create a heatmap and hierarchical clustering of the co-expression networks is performed using Euclidean distances of their similarities to pathways.

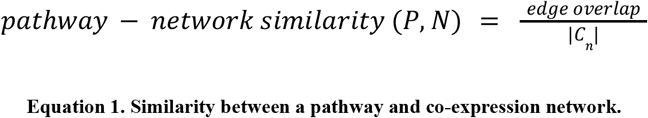

### 2.5. Implementation

Scripts to retrieve and process the datasets as well as to deploy the web application are available at https://github.com/ContNeXt. We have also provided comprehensive documentation to modify the filtering steps and add extensions to the scripts. For network analysis and visualizations, we used the Python NetworkX library (Hagberg *et al*., 2008) (https://networkx.github.io/), and Matplotlib, and seaborn, respectively. The processed data used in this work is available at Zenodo at https://zenodo.org/record/5831786.

## 3. Results

In **Section 3.1**, we provide an overview of the co-expression and PPI networks, while in **Sections 3.2-3.4**, we outline each of the analyses conducted, specifically at the protein-, network-, and pathway-levels **(Figure 2)**. Finally, **Section 3.5** presents ContNeXt, a web application developed to explore the results of this work.

**Figure 2.**
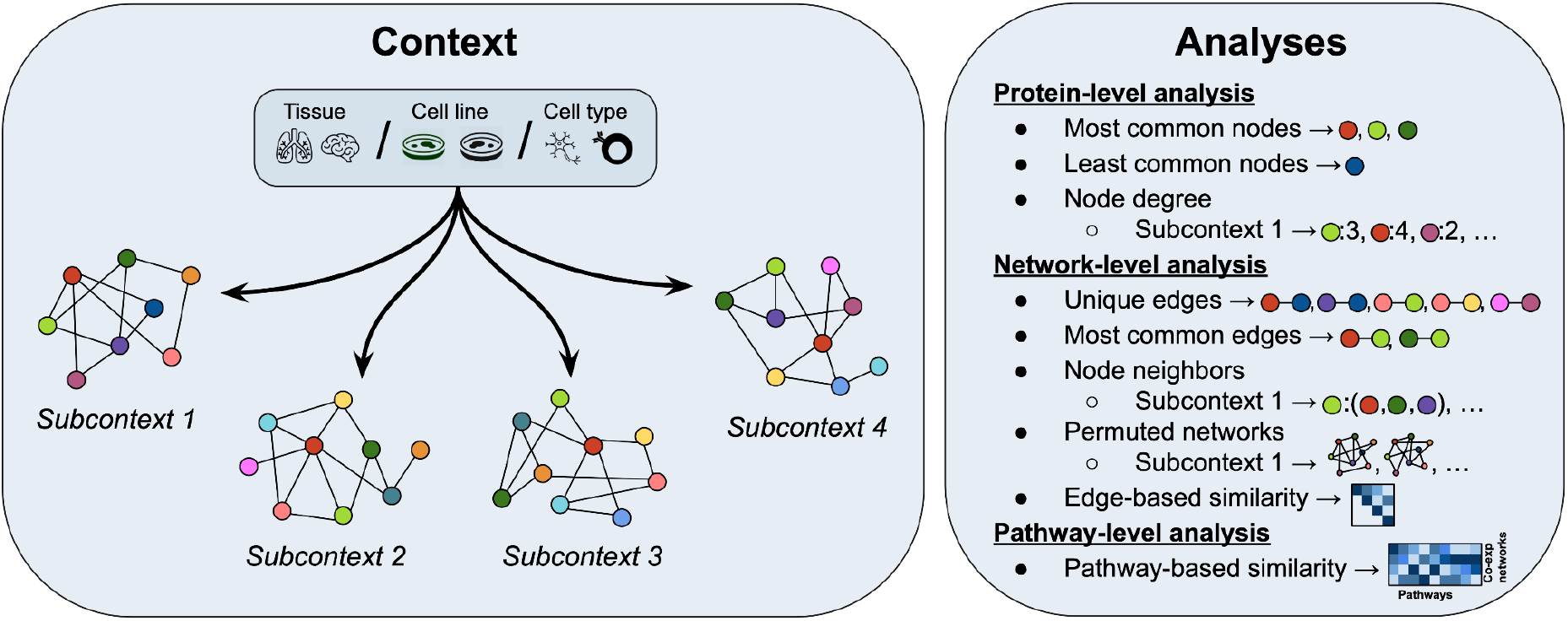
Overview of analyses conducted across all subcontexts in three different contexts (i.e., tissues, cell lines, and cell types). At the protein-level, patterns surrounding each single node are investigated (**Section 3.2**). The network-level analysis focuses on the relations between nodes (or node pairs) (**Section 3.3**) and the pathway-level analysis leverages defined node and edge sets to gain insights on context-specific co-expression networks (**Section 3.4**).

### 3.1. Overview of co-expression networks and interactome

From 364, 222, and 103 (at times overlapping) datasets that were categorized into 46 distinct tissues, 30 distinct cell types, 22 distinct cell lines, respectively, we systematically constructed co-expression networks corresponding to each of these contexts. **Figure 3** summarizes the size of each corresponding co-expression network. We find that across different contexts, the collected data, which depends on the study objectives, is biased towards certain groups of related subcontexts. For instance, in the tissue context, a large number of subcontexts belong to tissues of the nervous system, while in the cell type context, the majority of subcontexts are related to the immune system. This bias can especially be seen in the cell line context, where nearly all cell lines are derived from cancer cells. Finally, we investigated the correlation between the number of samples or datasets used to generate the co-expression networks and the size of the networks as a potential source of bias. We found no such dependency between the number of samples or datasets and the network size (**Supplementary Figure 1**).

**Figure 3.**
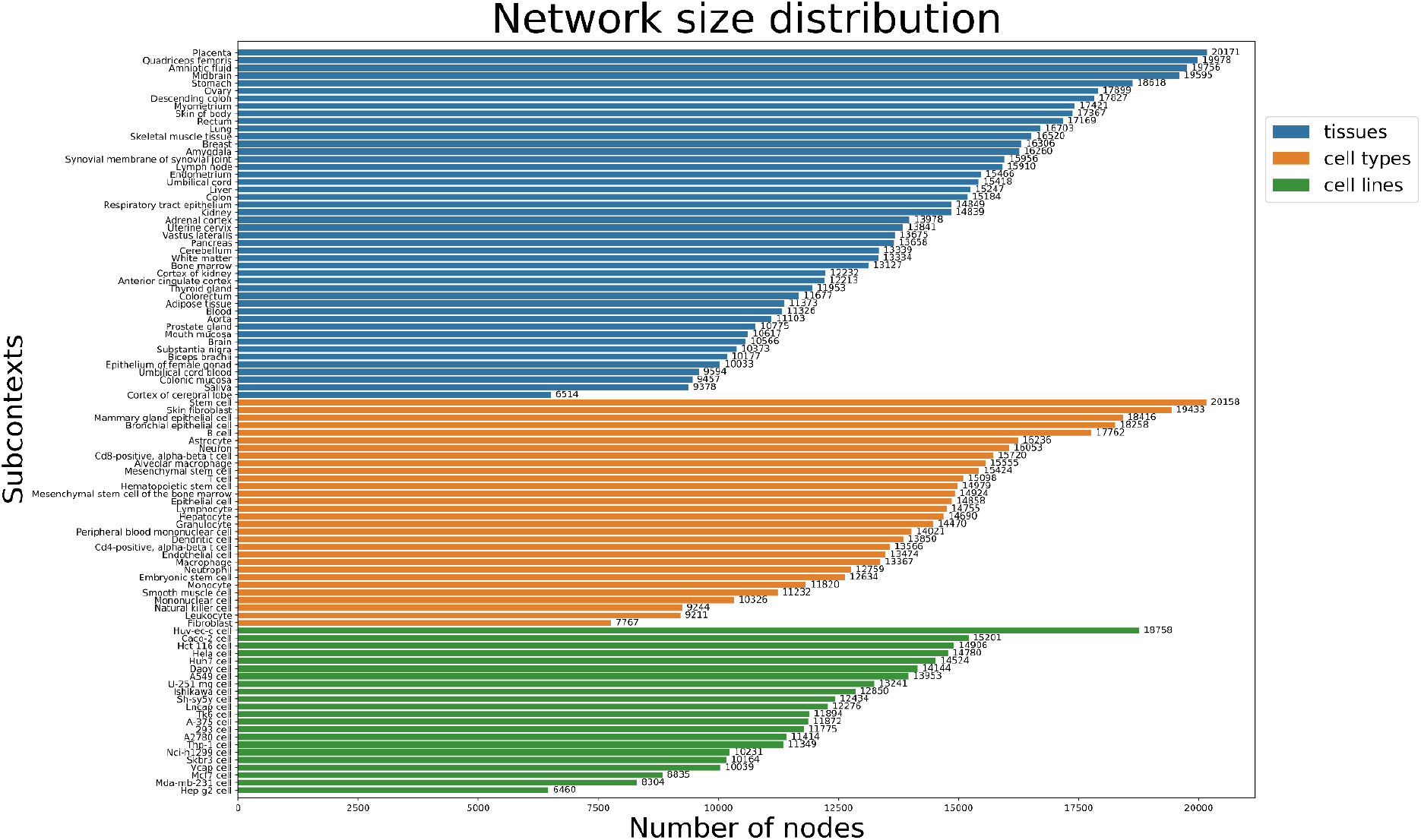
Distribution of network size for each of the three contexts. Distributions of network size are given as the number of nodes in each subcontext. In the tissue context, the cortex of cerebral lobe network had the fewest number of nodes (i.e., 6,514), while the placenta network had the largest number of nodes (i.e., 20,171) across not only all networks of the tissue context, but also across all other contexts. In the cell type context, the fibroblast network had the least number of nodes (i.e., 7,767), while the stem cell network had the highest number of nodes (i.e., 20,158). In the cell line context, the HepG2 cell line network had the least number of nodes (i.e., 6,460), while the Huv-ec-c cell line network had the largest number of nodes (i.e., 18,758). Generally, the networks within each context tended to vary greatly in size. For example, the tissue context includes networks ranging in size from 6,514 to 20,171 nodes.

With 8,601 nodes and 199,535 edges, although the size of the human interactome is on the same scale as other published human interactomes in recent studies (Luck *et al*., 2020; Vinayagama *et al*., 2016), the number of nodes (proteins) is less than half of the largest co-expression network. This was to be expected, as the majority of proteins measured in transcriptomic experiments have not yet been investigated in the literature and little is known of their functionality. Nodes of the interactome can be visualized in the web application **(see Section 3.5)** along with their neighbors, betweenness centrality, degree centrality, controllability classification, and information on whether the node is a housekeeping gene.

In order to discern unique features of context-specific co-expression networks which could be of biological significance, we first sought to identify genes known to arise from generic processes whose patterns are more likely to be stable and unaffected by any given context or condition. In particular, we investigated the presence of these, so called, housekeeping genes in each of the co-expression networks, noting that these genes are indicative of shared biology given their role in cell maintenance, and therefore, exhibit constant expression levels across all cells and conditions (Eisenberg and Levanon, 2003). Thus, by better understanding which genes have critical roles in basic cell maintenance, we could better direct our focus in determining genes of interest. The housekeeping genes dataset made available from Eisenberg and Levanon (2003) consisted of 3,804 genes (**Supplementary Table 4)**, 1,723 of which were present in the interactome (20% of the overall interactome).

To analyze the structural properties of the interactome, we employed an algorithm **(see Methods)** that has been applied to identify the importance of nodes and edges in biological networks **(Supplementary Text 1)**. The results of the controllability analysis indicate that the interactome has 1,233 driver nodes with which the network can be controlled. Overall, 74.6% of the nodes were classified as neutral, 16.17% dispensable, and 9.2% indispensable. A list of the full classifications can be seen in **Supplementary Table 5**, and a summary of these nodes can be seen in **Table 1**. We observed that the indispensable nodes were highly connected, as expected, had the highest average betweenness centrality, and a significant portion (i.e., ∼25%) were housekeeping genes. By comparison, neutral nodes were found to have half as many connections and an average betweenness centrality 10 times lower than indispensable nodes. However, the proportion of neutral nodes that were housekeeping genes were comparable to that of the indispensable nodes. By contrast, differences between the dispensable and indispensable nodes were far more pronounced; the average degree of dispensable nodes was only ∼6, compared to ∼107 for indispensable nodes, while the average betweenness centrality was more than 1,000 times lower. Additionally, only ∼8% of dispensable nodes were housekeeping genes, compared to roughly a quarter for both indispensable and neutral nodes.

**Table 1.**
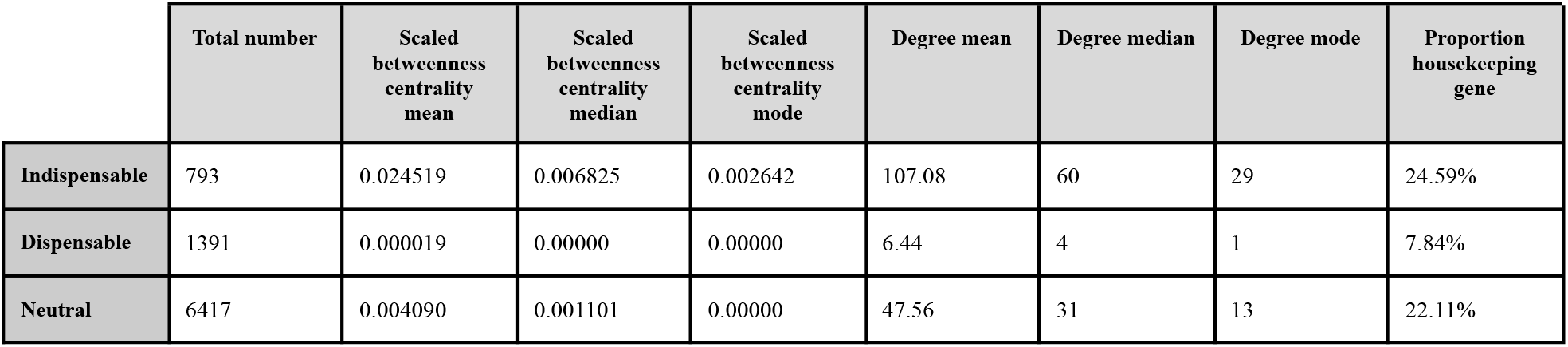
Regarding the interactome controllability, 6,417 of the total nodes (74.6%) were classified as neutral; i.e., removing them will have no effect on the number of driver nodes in the network, representing the largest proportion of nodes in the interactome. 1,391 (16.17% of the interactome) nodes were dispensable, meaning their removal would decrease the number of driver nodes in the network. Lastly, 793 nodes (9.2% of the interactome) were determined to be indispensable, which caused an increase in the need for driver nodes at their removal. In all three categories (i.e., betweenness centrality, degree, and housekeeping gene proportion), indispensable nodes had the highest value, followed by neutral, and dispensable with the lowest values. The indispensable nodes are listed in **Supplementary Table 6**. Betweenness centrality scores were scaled between 0 and 1 to facilitate comparability.

### 3.2. Analyses at the protein-level

We begin by exploring general trends for all co-expression networks of each context at the protein-level by focusing on the most and least common proteins (i.e., present in all or exactly one network within a context). We first used the results of the previously-mentioned controllability analysis of the interactome as well as housekeeping genes and overlapped them with the most and least common proteins in each context. As summarized in **Table 2**, of the most common nodes (i.e., proteins that could be found in each network within a given context), we found that the cell type context had the largest number of proteins across all networks (301 proteins), while the tissue network had the fewest (22 proteins). Among the most common nodes, the ratio of housekeeping genes was greater than the proportion of housekeeping genes present in the interactome (i.e., 20%), comprising nearly 50% of the most common nodes in each of the contexts.

**Table 2.** Most and least common proteins per context. The most and least common proteins of the co-expression networks (i.e., in all or exactly one network within a context) were overlapped with proteins given distinct classifications from the controllability analysis of the interactome as well as with housekeeping genes. 22 proteins were identified as the most common proteins, that is, found in all 46 co-expression networks of the tissue context. Of the 30 co-expression networks of the cell type context, 301 proteins were found in all of them, while among 22 co-expression networks in the cell line context, 185 proteins were identified in each network By comparison, no proteins were found to be unique to a single co-expression network in the tissue context, while only one was found in the cell type context belonging to the stem cell co-expression network. On the other hand, 106 least common proteins were found in the cell line context, only one of which is a housekeeping gene and none of which are indispensable. A full list of the proteins found in all or in a single network per context can be seen in **Supplementary Table 7**.

|  | Tissue context |  | Cell type context |  | Cell line context |  |
| --- | --- | --- | --- | --- | --- | --- |
| **Proteins in all co-expression networks** | **22** | 2 indispensable<br>11 housekeeping | **301** | 21 indispensable<br>180 housekeeping | **185** | 15 indispensable<br>81 housekeeping |
| **Proteins unique to one co-expression network** | **0** | 0 indispensable<br>0 housekeeping | **1** | 0 indispensable<br>0 housekeeping | **106** | 0 indispensable<br>1 housekeeping |

#### 3.2.1. Overlap of co-expression networks with the interactome

While only considering the proteins present in the interactome as well as at least one co-expression network, we conducted an in-depth investigation of whether proteins in the co-expression networks of a given context could consistently be identified in the human interactome network. We first noted trends at the protein-level by comparing the most and least common proteins across co-expression networks within a context against the most and least connected proteins of the interactome. As the co-expression network and interactome sizes vastly differed, we studied this overlap considering the top or bottom most proteins in proportions roughly equivalent in size. We selected various cut-offs for each context, corresponding to the number of co-expression networks (see **Supplementary Text 2 for** details on the cut-offs for each context). This ensured the inclusion of either the maximal or minimal possible overlap of the common proteins of the co-expression networks and connected proteins of the interactome, depending on whether our investigation focused on the most commonly or most uniquely occurring proteins, respectively. A detailed list of the resulting overlaps can be seen in **Supplementary Table 8**.

#### 3.2.2. Most common proteins

First, we focus on the most common proteins. Among the most commonly occurring proteins in the tissue context that overlapped with proteins from the interactome, a number of proteins belonged to the MAPK protein family (**Supplementary Table 8**). Proteins in this family are instrumental in transduction of extracellular signals to cellular responses and complex cellular processes such as apoptosis, development, differentiation, proliferation, and transformation (Zhang and Liu, 2002). While only the larger two comparisons in the tissue context **(Supplementary Figure 2;** lower two diagrams) resulted in an overlap, a significant portion of these overlapping proteins were also indispensable, or housekeeping. Within the large overlaps between the common cell type proteins and most connected interactome proteins (**Supplementary Figure 3**), a larger proportion of housekeeping genes was found than in any of the contexts studied, with more than half of each overlap being a housekeeping gene (i.e., 50-67%), and more of the proteins are also indispensable.

In cell lines, we observed a substantial overlap of most common proteins that are also found in the interactome overall, including when using the strictest cut-offs, however, significantly less were found to be indispensable or housekeeping than in the tissue and cell type contexts (**Supplementary Figure 4**). We select a proportional set from each context (400 of the most common proteins per context) to compare their overlaps with the interactome (**Supplementary Table 9A**). The overlaps all had a similar number of proteins in them, between 30 and 37 proteins. Across contexts, there was a similar proportion of the overlap which are indispensable nodes of the interactome (∼32% in tissues, 40% in cell types, and ∼43% in cell lines). On the other hand, the proportion of housekeeping genes varied more, with 43% of the proteins from the cell line overlap, while tissues and cell types both had more than 60%. Overall, housekeeping genes seem to be best represented in the co-expression networks. We observed a number of proteins in all of the context’s overlaps belonging to the Ribosomal protein (RP) family (**Supplementary Table 9A**), from both small and large subunits. RPs are essential in protein synthesis (Yoshihama *et al*., 2002). The tissue overlap had one from large and one from small subunit, the cell type overlap had four from large and one from small subunit, and the cell line overlap had one small subunit RP. We also found that the average number of relations for the proteins in the interactome that overlapped with the approximately top 400 most common proteins in the tissue and cell line networks (∼73 and ∼72 relations, respectively), was much higher than the average number of relations overall in the interactome (∼46 relations). This suggests that the common tissue- and cell line-wide proteins across the co-expression networks are better represented in the scientific literature. In the cell type networks, this average was less high, ∼60 relations, but still more than overall in the interactome.

#### 3.2.3. Least common proteins

Next, we investigated the least common proteins in the co-expression networks and their overlap with the least connected proteins in the interactome. This time, the tissue context presented a more consistent overlap while increasing the protein pool, but still a minimal overlap (**Supplementary Figure 5**). The overlap with the interactome and the cell type context was about the same as in the tissue context (**Supplementary Figure 6**). In the cell line context, we found a small, steadily increasing overlap with each interval comparison, which was not the case in the most common proteins (**Supplementary Figure 7**). The overlap with the interactome in the larger comparisons was roughly the same as in every other context. The minimal overlaps suggest that little is currently known of these proteins. Additionally, we also selected proportional sets of the 400 least common proteins in each context, also occurring the interactome overall against the 400 least connected nodes of the interactome (**Supplementary Table 9B**). The sizes of the overlap didn’t vary as much as in the most common and connected comparison, with each context having around 30 proteins in the overlap. As expected, with these overlaps, either one or no proteins are also indispensable or housekeeping. We observe an overwhelming number of proteins belonging to the ZNF protein family in each of the overlaps (i.e., 10/34 (29%) in tissues, 11/33 (33%) in cell types, and 4/27 (15%) in cell lines) (**Supplementary Table 9B**). While ZNFs are widely found in the organism, they play critical roles in specific tissues, and in the development of many diseases (Cassandri *et al*., 2017).

### 3.3. Analyses at the network-level

We first focused on analyzing edges of the co-expression networks, including the unique and most commonly occurring edges within contexts. Additionally, we leveraged prior knowledge from a referential human interactome and studied the correspondence of edges from this network against the strongest pairwise correlations of the co-expression networks. Subsequently, we validated these findings by conducting an equivalent comparison against randomly generated versions of the co-expression networks. Finally, we conducted a similarity analysis on the network edges within each context.

#### 3.3.1. Unique and most commonly occurring edges

We first assessed whether there were any edges specific to particular tissue networks, identifying 45,963,343 unique edges in total (i.e., 49% of all edges). We also identified 34,584,720 unique edges in the cell type context (i.e., 57% of all edges) and 31,941,789 unique edges in the cell line context (i.e., 71% of all edges). These proportions are similar to findings by Stacey *et al*. (2018) who found that over half of edges in several PPI databases are context-specific. **Figure 4** illustrates the frequency of unique and common edges in all networks within a context. We find that edges which are common to at least 25% of networks within a context are rare (i.e., between 0.07% and 0.16%), while those which are in at least 75% of networks are nearly negligible (i.e., 33 edges in total for tissues, 9 for cell types, and 4 for cell lines). As only the 1% strongest correlations were selected for each network, it was foreseen that a large number of edges in our resulting co-expression networks would be specific to a single subcontext. Although these unique edges are interesting to explore for a given subcontext (green portions in **Figure 4**), given the sheer volume of unique edges, their investigation was outside of the scope of this work.

**Figure 4.**
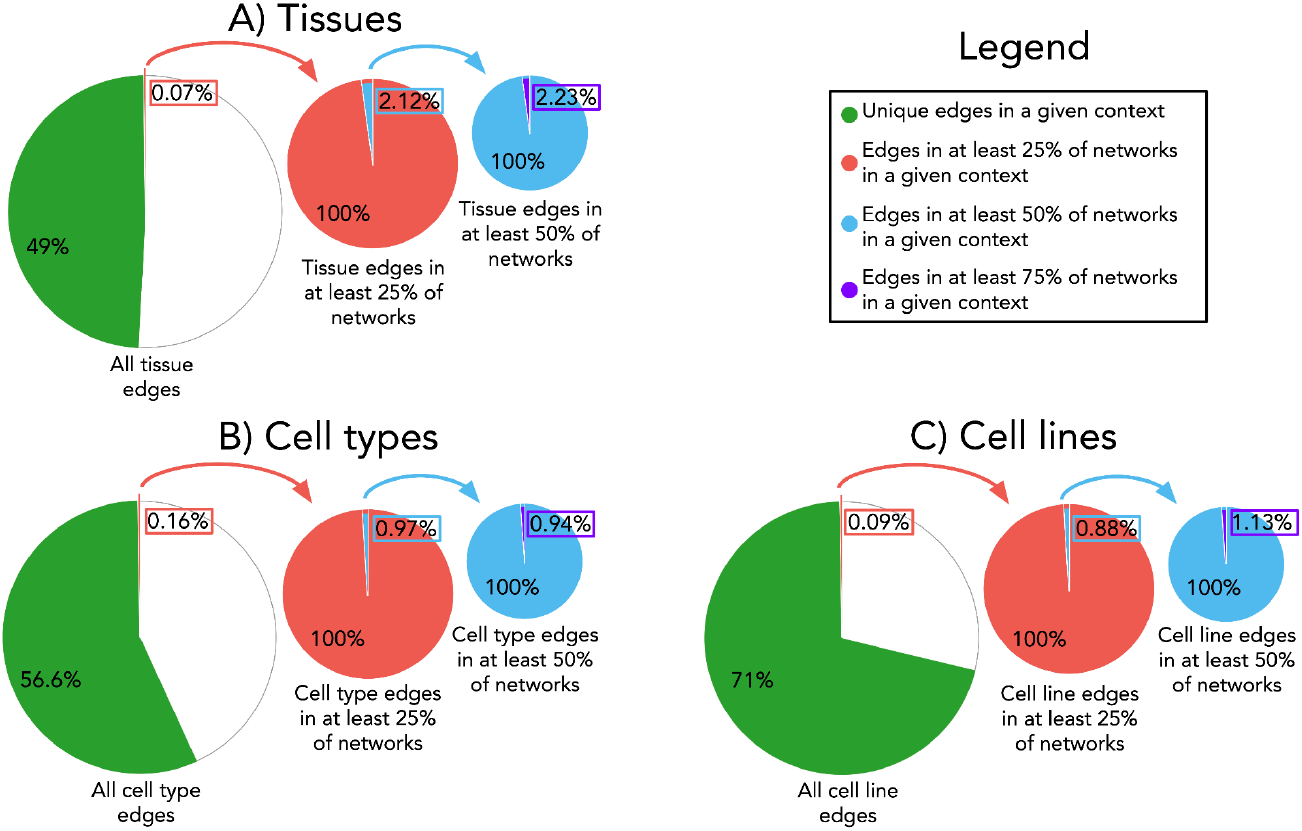
Frequency of edge occurrence across networks within a context. Proportions of edges are given as those that are unique, or common to varying degrees, in networks within the **A)** tissue, **B)** cell type, and **C)** cell line context. From the total set of edges that occur across all networks within each context, the fraction of edges that are unique (i.e., appear in at most one network within a given context) are shown in green. From this total set of edges, the fraction of those which appear in at least 25% of networks within a given context are magnified in a consecutively smaller pie chart (i.e., predominantly in red). Similarly, those which appear in at least 50% of networks within a given context are magnified and illustrated in a pie chart predominantly in blue. Finally, of this latter group of edges, the fraction of edges that are most common (i.e., appear in at least 75% of all networks within a given context) are highlighted in purple.

We hypothesize that these common edges correspond to basal correlations that are not specific as they appear in the majority of networks within one or more contexts. Thus, we analyze the most frequently occurring edges in each of the three contexts. Unsurprisingly, the two housekeeping genes of the tubulin alpha families (i.e., TUBA1C and TUBA1B) are nearly always found to be connected to each other (in 83 out of 98 networks), regardless of context. Additionally, IFITM2 and IFITM3, proteins of the interferon-induced transmembrane family, which play a key role in immune system functions, are also often seen connected to each other in 84 out of 98 networks. Members of the human leukocyte antigens (HLA) protein family are also often interconnected across the cell type and cell line contexts. This is in line with Crow *et al*. (2019) who found that certain gene modules are predictably found across biological conditions, such as those of the immune response. In our previous paper (Figueiredo *et al*., 2021), we found that of the most common edges among 63 major diseases, members of the Metallothionein (MT) family of proteins, were in nearly half of these edges. Similarly, here again we observed that a large number of MT proteins share neighbors across networks in every context.

Of the most common edges throughout all contexts **(see Supplementary Text 3**), none were indispensable within the interactome. When widening our search to the top 100,000, we found only seven, three, and one edge in the tissue, cell type, and cell line contexts to be indispensable in the interactome, respectively. Next, these most common edges found in the majority of networks of a given context were compared to the interactome network to identify concordance between the two. We performed a range of comparisons on the most common edges by focusing only on the top 1,000 to 10,000 edges, in increments of 1,000. Then, the most common edges in each co-expression network were compared to the interactome. Overall, we found little overlap in the most common edges. In the tissue context, we found an overlap of only 5% in the top 1,000 most common edges against the interactome, with this overlap decreasing to 4% when considering the top 10,000 most common edges. In comparison these proportions ranged from ∼7% to 3% in the cell type context between the top 1,000 and 10,000 most common edges, and 4% to 2% in the cell line context.

#### 3.3.2. The strongest correlations tend to correspond with protein-protein interactions more than expected by chance

In this section, we investigate whether the strongest correlations present in the co-expression networks correspond to PPIs more often than what would be expected by chance. For this purpose, we permuted each co-expression network for each context 1,000 times while maintaining the original graph structure **(see Methods)**. We next compared the overlap of edges between these permuted co-expression networks with the human interactome (the results of the first 100 permutations can be seen in **Supplementary Table 10)**. Our results show that, on average, the original co-expression networks have 1.55 times as many edges in common with the human interactome as compared to the permuted networks, which exhibited a comparatively low variability in their overlap within a subcontext. Across all contexts, the maximum difference in overlap was for the ovary subcontext, where the original ovary co-expression network had 3.3 times as many edges in common with the interactome as compared to the permuted versions. In comparison, the saliva co-expression network showed the smallest difference in edge overlap between the original and permuted co-expression networks, with the overlap of the interactome with the original co-expression network having only 1.01 times as many edges as the permuted versions on average. Thus, we find that co-expression patterns correspond with PPIs more than expected by chance.

#### 3.3.3. Edge-based similarity across co-expression networks

Next, we investigated edge similarity across networks within a given context. By comparing the co-expression networks to each other rather than just the interactome, we could identify the networks that were most similar edgewise. In the tissue context, two pairs of networks displayed the highest degree of similarity, namely the brain and the cortex of the cerebral lobe, and the colon and the rectum (**Figure 5A**). This finding was not surprising given that these pairs of tissues are anatomically related (i.e., both are of the brain or the colorectum). The cell line context had a few standout pairs of networks which had the highest degree of similarity (**Figure 5B**). Specifically, the highest similarity was between two different human breast cancer cell lines: MDA-MB-231 and MCF7. Additionally, the MCF7 cell line again had a high similarity with a human colon cancer cell line, HCT 116. On the other hand, in the cell type context, rather than specific pairs showing the highest similarity with each other, a few selected subcontexts had a high similarity with most of the other networks overall (**Figure 5C**). These include the macrophage and peripheral blood mononuclear cell networks, which had high similarity with more than half of the other networks.

**Figure 5.**
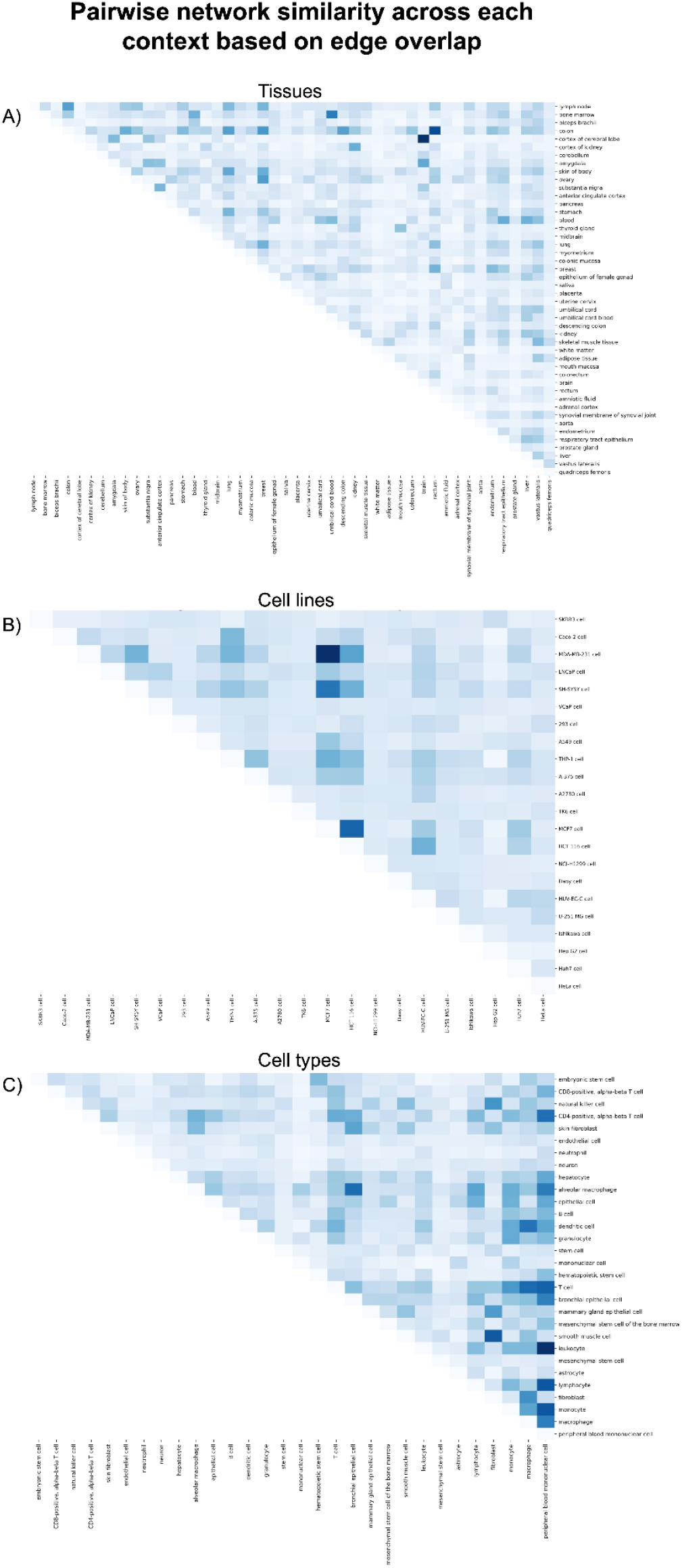
Pairwise co-expression network similarity across contexts. For each pair of co-expression networks within a given context, edge overlap was calculated as a measure of similarity between networks for the **A)** tissue, **B)** cell line, and **C)** cell type contexts. A high quality version of the figure is available at https://github.com/ContNeXt/scripts/blob/main/figures/figure5.pdf.

### 3.4. Mapping co-expression networks to pathway knowledge

Lastly, we attempted to establish patterns across co-expression networks at a pathway-level by overlaying pathway knowledge with the co-expression networks. If a given pathway is related to a specific network (e.g., fatty acid metabolism pathway and the liver co-expression network), we would expect that the proteins in the pathway would be strongly correlated in the co-expression network. Furthermore, we assume that, given a set of highly co-expressed genes of which a majority are involved in a particular pathway, the remaining genes may be functionally relevant to the pathway as well. We therefore seek to identify the pathways associated with networks from each of the investigated contexts. Using the KEGG database (Kanehisa *et al*., 2021), we mapped pathway knowledge to co-expression networks according to **Equation 1 (see Methods)**.

We found several groups of tissues that had high similarities with pathways related to the given tissues **(Figure 6)**. For instance, the two tissue networks corresponding to cortex of cerebral lobe and brain shared a large group of pathways exhibiting a high degree of similarity, including nine synaptic pathways **(Figure 6; green oval) (Supplementary Table 11)**. Furthermore, the three networks for liver, cortex of kidney, and kidney also had the highest level of similarity with numerous pathways, including eight involving the regulation of fatty acids as well as 11 involving amino acid metabolism and degradation (**Figure 6; red oval)** (**Supplementary Table 12**). Not surprisingly, the adipose tissue network also showed the highest similarity with adipose-related pathways, such as adipocytokine signaling pathway and regulation of lipolysis in adipocytes pathway.

**Figure 6.**
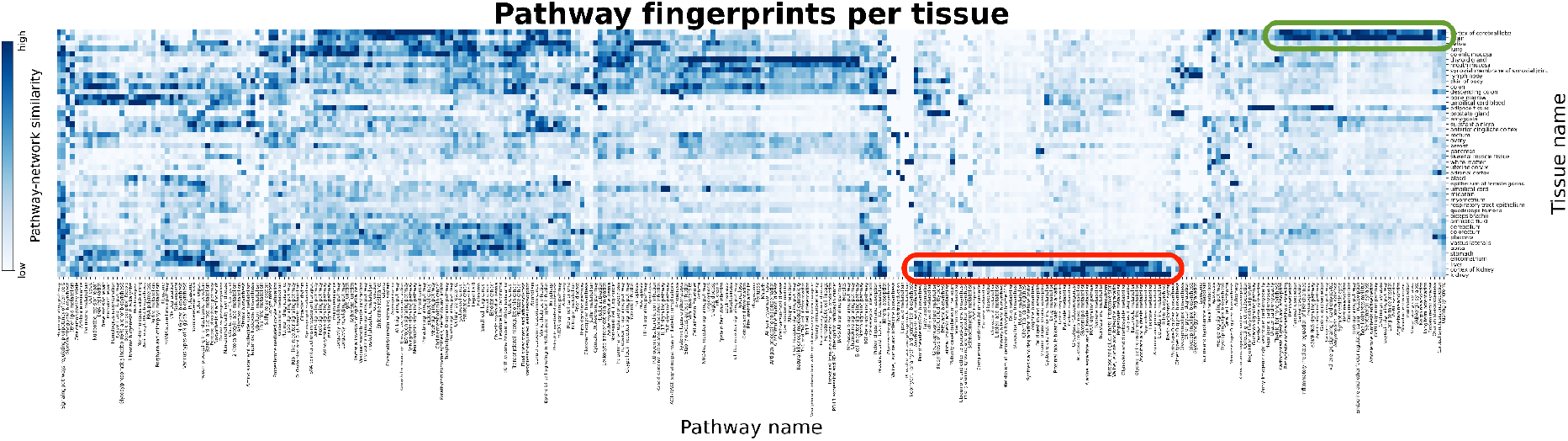
Similarity between tissue-specific co-expression networks and KEGG pathways. The similarity between a particular pathway and a co-expression network is defined as the percentage of pairwise combinations of proteins of a given KEGG pathway that can be found in a co-expression network as edges. Light blue corresponds to a lower similarity, while dark blue corresponds to a high similarity. A high quality version of this figure is available at https://github.com/ContNeXt/scripts/blob/main/figures/figure6_highquality.pdf and can also be visualized in the web application.

In the cell type context, while no groups of network shared distinct pathways among them, we found three cell types having distinct groups of pathways with very high similarity unique to a single network. For example, a number of pathways showed a high degree of similarity to the neutrophil co-expression network **(Supplementary Figure 8; red oval)**, namely, 11 that regulate the immune response (**Supplementary Table 13**). Additionally, the co-expression network for hepatocytes, the primary cell type of the liver, had the highest level of similarity with many pathways **(Supplementary Figure 8; yellow oval)**, including six involving basic liver function as well as many metabolic pathways, particularly 10 pertaining to amino acids metabolism and seven for other specific molecules (**Supplementary Table 14**). Lastly, we found an additional group of pathways that were exclusively similar to one network, namely the neuron (**Supplementary Figure 8; green oval)**. Specifically, this included five pathways related to neurotransmitter systems, long-term depression, and pathways related to addiction (**Supplementary Table 15**).

Analogous to the cell type context, while related groups of networks from the cell line context were not found to be similar to related groups of pathways **(Supplementary Figure 9)**, several individual cell lines were observed to be highly similar to a group of pathways. However, these pathways were not necessarily unique to the cell line, showing some similarity with other cell lines as well. Interestingly, we found a large group of pathways (i.e., 70 in total) with consistently high similarity with nearly all cell lines, with the exception of the THP-1 cell line **(Supplementary Figure 9; green rectangle)**. These include 24 different signaling pathways and 16 different cancer pathways **(Supplementary Table 16)**. Notably, we found a group of pathways that were distinctly similar to two cell lines (i.e., A549 and TK6). Specifically, 14 pathways showed a high degree of similarity to the A549 cell line co-expression network **(Supplementary Figure 9; yellow oval)**. This cell line originated from adenocarcinomic human alveolar basal epithelial cells from lung cancer and is used as a model for drug metabolism (Foster *et al*., 1998). Of these 14 pathways that, on average, showed the highest similarity to this cell line relative to the others, eight were pathways involving metabolism and three were pathways related to compound biosynthesis (**Supplementary Table 17**). Similarly, we identified a group of pathways which showed a higher similarity to the TK6 cell line, originating from a human B lymphoblastoid cell (Schwartz *et al*., 2004), over all other cell lines **(Supplementary Figure 11; red oval)**, including five signaling pathways (**Supplementary Table 18**).

### 3.5. ContNeXt – a web application to explore gene expression patterns across contexts

To provide access to the co-expression networks and analyses presented in this work, we have developed ContNeXt, a web application that facilitates the large-scale exploration and analysis of transcriptomic patterns across multiple contexts. The main page of the web application allows users to search co-expression patterns for a given node in a particular context or browse and query specific nodes in a certain subcontext (**Figure 7A**). With interactive network visualizations, users can explore these patterns and employ functionalities such as filtering or search boxes **(Figure 7B)**. Similarly, the heatmaps presented in this work can be interactively explored through the web application **(Figure 7C)**. Finally, both the processed data and networks can be downloaded directly from the web application.

**Figure 7.**
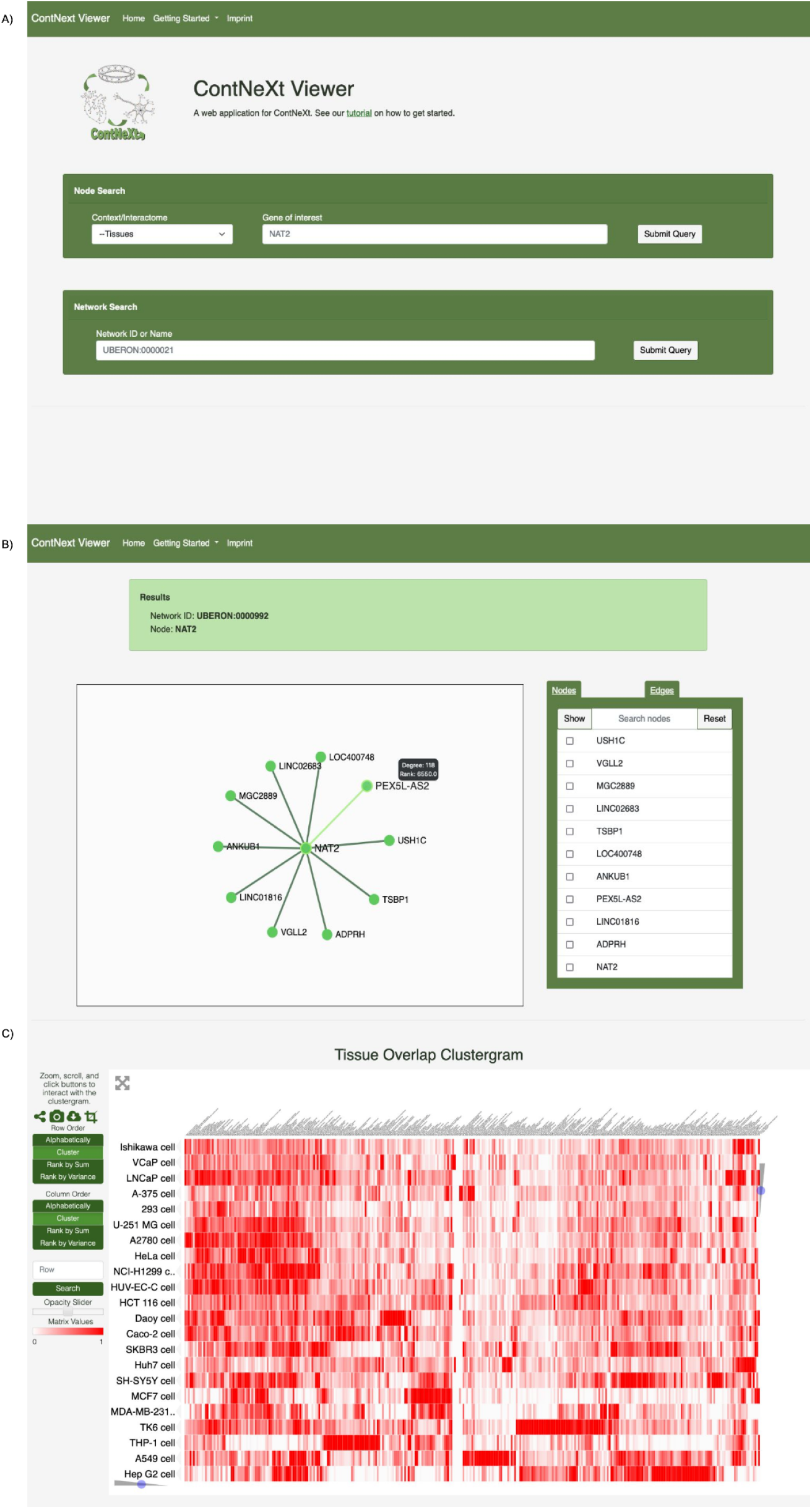
ContNeXt web application. A) Main page. Users can query for specific genes or directly explore the networks of a given context. B) Network page. Users can explore and navigate through the neighbors of a specific gene for each network. C) Heatmap visualization. Heatmaps presented in this work can be interactively viewed to investigate pairwise co-expression network -based similarity as well as pathway-co-expression network -based similarity.

## 4. Discussion

We have presented a large-scale network-based approach that aims at revealing common and specific biological processes and mechanisms across contexts by identifying transcriptional patterns that are unique to various cell types, tissues, and cell lines, as well as patterns which are consistent across them. In order to do so, we constructed co-expression networks to capture the strongest correlations observed in 98 specific subcontexts belonging to these three biological contexts (i.e., tissues, cell types, and cell lines) and conducted a series of analyses at the protein, network, and pathway levels. Finally, we developed a web application to enable users to query and display these networks and ultimately, explore shared and distinct co-expression patterns for multiple contexts.

We believe that one strength of our work is its robustness, as we have systematically leveraged hundreds of curated datasets, thereby ensuring a diverse sample of experiments conducted in similar settings whilst applying a common preprocessing and analysis pipeline. However, although we applied a conservative inclusion/exclusion criteria, we cannot assume that every dataset in the same (sub)context is equivalent and thus, some of the patterns identified may be dataset-specific. To account for this factor and reduce noise and variability across datasets, we focused on the 1% strongest correlations, keeping in mind that the choice of cut-off can influence the resulting co-expression network (Yip and Horvath, 2007), and also constrained our analysis to subcontexts with a large number of samples. Still, independently of this minimum criteria, there are differences in the number of datasets per subcontext that could lead to variability for specific subcontexts with a small sample size. Another limitation is that we have exclusively relied on the platform with a large number of datasets in the Gemma database. Similarly, we also employed Gemma’s context annotations to classify the datasets. While it is technically possible to include more platforms in our analysis as well as annotate datasets from other databases, each additional platform would require its own independent processing pipeline and a significant curation effort. Furthermore, in the cell line context, it is important to note that the majority of cell lines originate from widely used immortal cancer cell lines, which might differ from the normal human cells used for the cell type and tissue contexts. Finally, we would like to remark on two other limitations of our analysis. Firstly, while we employed a large and high-quality version of the protein-protein human interactome, some parts of the graphs are more dense than others as some proteins are under-studied (Schaefer *et al*., 2015). Secondly, some of the analyses are influenced by the size of the co-expression networks **(Figure 3)**, as the fewer nodes a network has, the more dense it is due to the larger amount of connections between its nodes.

Lastly, we would like to mention some of the prospects we foresee for future work. Firstly, by shifting the analysis towards single-cell experiment datasets, we can potentially identify more granular patterns. Furthermore, single-cell RNA-seq data can be used to verify whether the observed tissue-specific transcriptional patterns are indeed characteristic to specific tissues, or are influenced by their cellular composition, as observed by Farahbod and Pavlidis (2020). While this large-scale exercise is not feasible at the moment due to the lack of available data of this kind, we expect that it could be conducted in future. Secondly, disease-specific gene expression datasets can be exploited to compare disease-specific signatures with the ones observed in a related normal tissue or cell type in order to identify the biological processes and pathways that are dysregulated in the disease context. Thirdly, as demonstrated by Azevedo *et al*. (2021) and Sealfon *et al*. (2021), machine learning models could be trained on the generated co-expression networks to classify signatures coming from new samples into a particular context given its specific characteristics.

## Data availability

All data supporting the conclusions of this article are available at https://zenodo.org/record/5831786 and scripts can be found at https://github.com/ContNeXt/scripts. ContNeXt and its source code are available at https://contnext.scai.fraunhofer.de and https://github.com/ContNeXt/web_app, respectively.

## Authors’ Contributions

DDF and SM conceived and designed the study. RQF and TR processed the transcriptomic datasets. RQF implemented the methodology and analyzed the results supervised by SM and DDF. SDS implemented the web application. ATK, MHA and DDF acquired the funding. RQF, SM, and DDF wrote the manuscript. ATK and MHA reviewed the manuscript.

## Acknowledgements

We would like to thank the entire Gemma team, especially Paul Pavlidis, for their support using their tool. Furthermore, we would like to thank André Gemünd for his technical assistance.

## References

1. Azevedo T, Dimitri GM, Lió P, and Gamazon ER. (2021). Multilayer modelling of the human transcriptome and biological mechanisms of complex diseases and traits. NPJ systems biology and applications, 7(1), 1–13. https://doi.org/10.1038/s41540-021-00186-6

2. Cassandri M, Smirnov A, Novelli F, Pitolli C, Agostini M, Malewicz M, et al. (2017). Zinc-finger proteins in health and disease. Cell death discovery, 3(1), 1–12. https://doi.org/10.1038/cddiscovery.2017.71

3. Crow, M., Lim, N., Ballouz, S., Pavlidis, P., and Gillis, J. (2019). Predictability of human differential gene expression. Proceedings of the National Academy of Sciences, 116(13), 6491–6500. https://doi.org/10.1073/pnas.1802973116

4. Diehl AD, Meehan TF, Bradford YM, Brush MH, Dahdul WM, Dougall DS, et al. (2016). The Cell Ontology 2016: enhanced content, modularization, and ontology interoperability. Journal of biomedical semantics, 7(1), 1–10. https://doi.org/10.1186/s13326-016-0088-7

5. Dobrin R, Zhu J, Molony C, Argman C, Parrish ML, Carlson S, Allan MF, Pomp D, and Schadt EE. (2009) Multi-tissue coexpression networks reveal unexpected subnetworks associated with disease. Genome biology, 10(5), 1–3. https://doi.org/10.1186/gb-2009-105-r55

6. Edgar R, Domrachev M, and Lash AE. (2002). Gene Expression Omnibus: NCBI gene expression and hybridization array data repository. Nucleic acids research, 30(1), 207–210. https://doi.org/10.1093/nar/30.1.207

7. Eisenberg E., and Levanon EY (2013). Human housekeeping genes, revisited. TRENDS in Genetics, 29(10), 569–574. https://doi.org/10.1016/j.tig.2013.05.010

8. Farahbod M, and Pavlidis P. (2020). Untangling the effects of cellular composition on coexpression analysis. Genome research, 30(6), 849–859. https://doi.org/10.1101/gr.256735.119

9. Figueiredo RQ, Raschka T, Kodamullil AT, Hofmann-Apitius M, Mubeen S, and Domingo-Fernández D (2021). Towards a global investigation of transcriptomic signatures through co-expression networks and pathway knowledge for the identification of disease mechanisms. Nucleic acids research, 49(14), 7939–7953. http://doi.org/10.1093/nar/gkab556

10. Foster KA, Oster CG, Mayer MM, Avery ML, and Audus KL. (1998). Characterization of the A549 cell line as a type II pulmonary epithelial cell model for drug metabolism. Experimental cell research, 243(2), 359–366. https://doi.org/10.1006/excr.1998.4172

11. Hagberg AA, Schult DA, and Swart PJ. (2008). Exploring network structure, dynamics, and function using NetworkX. Proceedings of the 7th Python in Science Conference (SciPy2008), 11–15.

12. Hanhijärvi S, Garriga, GC, and Puolamäki K. (2009). Randomization techniques for graphs. In Proceedings of the 2009 SIAM International Conference on Data Mining (pp. 780–791). https://doi.org/10.1137/1.9781611972795.67

13. Jassal B, Matthews L, Viteri G, Gong C, Lorente P, Fabregat A, … and D’Eustachio P. (2020). The reactome pathway knowledgebase. Nucleic acids research, 48(D1), D498–D503. https://doi.org/10.1093/nar/gkz1031

14. Johnson WE, Li C, Rabinovic A. (2007). Adjusting batch effects in microarray expression data using empirical Bayes methods. Biostatistics, 8(1), [118-127. https://doi.org/10.1093/biostatistics/kxj037

15. Kanehisa M, Furumichi M, Sato Y, Ishiguro-Watanabe M, and Tanabe M. (2021). KEGG: integrating viruses and cellular organisms. Nucleic acids research, 49(D1), D545–D551. https://doi.org/10.1093/nar/gkaa970

16. Kitsak M, Sharma A, Menche J, Guney E, Ghiassian SD, Loscalzo J, and Barabási AL (2016). Tissue specificity of human disease module. Scientific reports, 6(1), 1–12. https://doi.org/10.1038/srep35241

17. Koussounadis A, Langdon SP, Um IH, Harrison DJ, and Smith VA (2015). Relationship between differentially expressed mRNA and mRNA-protein correlations in a xenograft model system. Scientific reports, 5(1), 1–9. https://doi.org/10.1038/srep10775

18. Langfelder P, and Horvath S. (2008). WGCNA: an R package for weighted correlation network analysis. BMC bioinformatics, 9(1), 1–13. https://doi.org/10.1186/1471-2105-9-559

19. Lee YF, Lee CY, Lai LC, Tsai MH, Lu TP, and Chuang EY. (2018). CellExpress: a comprehensive microarray-based cancer cell line and clinical sample gene expression analysis online system. Database, 2018. https://doi.org/10.1093/database/bax101

20. Lee J, Shah M, Ballouz S, Crow M, and Gillis J (2020). CoCoCoNet: conserved and comparative co-expression across a diverse set of species. Nucleic acids research, 48(W1), W566–71. https://doi.org/10.1093/nar/gkaa348

21. Lim N, Tesar S, Belmadani M, Poirier-Morency G, Mancarci BO, Sicherman J, et al. (2021). Curation of over 10,000 transcriptomic studies to enable data reuse. Database (2021), baab006. https://doi.org/10.1093/database/baab006

22. Liu YY, Slotine JJ, and Barabási AL. (2011). Controllability of complex networks. Nature, 473(7346), 167–173. https://doi.org/10.1038/nature10011

23. Lonsdale J, Thomas J, Salvatore M, Phillips R, Lo E, Shad S, et al. (2013). The genotype-tissue expression (GTEx) project. Nature genetics, 45(6), 580–585. https://doi.org/10.1038/ng.2653

24. Luck K, Kim DK, Lambourne L, Spirohn K, Begg BE, Bian W, et al. (2020). A reference map of the human binary protein interactome. Nature, 580(7803), 402–408. https://doi.org/10.1038/s41586-020-2188-x

25. McKenzie AT, Wang M, Hauberg ME, Fullard JF, Kozlenkov A, Keenan A, et al. (2018). Brain cell type specific gene expression and co-expression network architectures. Scientific reports, 8(1), 1–9. https://doi.org/10.1038/s41598-018-27293-5

26. Mungall CJ, Torniai C, Gkoutos GV, Lewis SE, and Haendel MA. (2012). Uberon, an integrative multi-species anatomy ontology. Genome biology, 13(1), 1–20. https://doi.org/10.1186/gb-2012-131-r5

27. Nusinow DP, Szpyt J, Ghandi M, Rose CM, McDonald III, E., Kalocsay M, et al. (2020). Quantitative proteomics of the cancer cell line encyclopedia. Cell, 180(2), 387–402. https://doi.org/10.1016/j.cell.2019.12.023

28. Obayashi T, Kagaya Y, Aoki Y, Tadaka S, and Kinoshita K. (2019). COXPRESdb v7: a gene coexpression database for 11 animal species supported by 23 coexpression platforms for technical evaluation and evolutionary inference. Nucleic acids research, 47(D1), D55–62. https://doi.org/10.1093/nar/gky1155

29. Oldham MC, Konopka G, Iwamoto K, Langfelder P, Kato T, Horvath S, and Geschwind DH. (2008). Functional organization of the transcriptome in human brain. Nature neuroscience, 11(11), 1271–82. https://doi.org/10.1038/nn.2207

30. Perkins AD, and Langston MA. (2009). Threshold selection in gene co-expression networks using spectral graph theory techniques. BMC bioinformatics 10(11), 1–11. https://doi.org/10.1186/1471-2105-10-S11-S4

31. Pierson E, GTEx Consortium, Koller D, Battle A, and Mostafavi S. (2015) Sharing and specificity of co-expression networks across 35 human tissues. PLoS computational biology, 11(5), e1004220. https://doi.org/10.1371/journal.pcbi.1004220

32. Rachlin J, Cohen DD, Cantor C, and Kasif S. (2006). Biological context networks: a mosaic view of the interactome. Molecular Systems Biology, 2(1), 66. https://doi.org/10.1038/msb4100103

33. Romero IG, Ruvinsky I, and Gilad Y. (2012). Comparative studies of gene expression and the evolution of gene regulation. Nature Reviews Genetics, 13(7), 505–516. https://doi.org/10.1038/nrg3229

34. Rung J, and Brazma A. (2013). Reuse of public genome-wide gene expression data. Nature Reviews Genetics, 14(2), 89–99. https://doi.org/10.1038/nrg3394

35. Sarntivijai S, Lin Y, Xiang Z, Meehan TF, Diehl AD, Vempati UD, et al. (2014). CLO: the cell line ontology. Journal of biomedical semantics, 5(1), 1–10. https://doi.org/10.1186/2041-1480-5-37

36. Schaefer MH, Serrano L, and Andrade-Navarro MA (2015). Correcting for the study bias associated with protein–protein interaction measurements reveals differences between protein degree distributions from different cancer types. Frontiers in genetics, 6, 260. https://doi.org/10.3389/fgene.2015.00260

37. Schwartz JL, Jordan R, Evans HH, Lenarczyk M, and Liber HL. (2004). Baseline levels of chromosome instability in the human lymphoblastoid cell TK6. Mutagenesis, 19(6), 477–482. https://doi.org/10.1093/mutage/geh060

38. Sealfon RS, Wong AK, and Troyanskaya OG (2021). Machine learning methods to model multicellular complexity and tissue specificity. Nature Reviews Materials, 1–13. https://doi.org/10.1038/s41578-021-00339-3

39. Sonawane AR, et al. (2017). Understanding tissue-specific gene regulation. Cell reports, 21(4), 1077–1088. https://doi.org/10.1016/j.celrep.2017.10.001

40. Stacey RG, Skinnider MA, Chik, JHL, and Foster, LJ. (2018). Context-specific interactions in literature-curated protein interaction databases. BMC genomics, 19(1), 1–10. https://doi.org/10.1186/s12864-018-5139-2

41. Trapotsi MA, Hosseini-Gerami L, and Bender A (2022). Computational analyses of mechanism of action (MoA): data, methods and integration. RSC Chemical Biology. https://doi.org/10.1039/D1CB00069A

42. The Gene Ontology Consortium (2021). The Gene Ontology resource: enriching a GOld mine. Nucleic Acids Research, 49(D1), D325–D334. https://doi.org/10.1093/nar/gkaa1113

43. Vinayagam A, Gibson TE, Lee HJ, Yilmazel B, Roesel C, Hu Y, et al. (2016). Controllability analysis of the directed human protein interaction network identifies disease genes and drug targets. Proceedings of the National Academy of Sciences, 113(18), 4976–4981. https://doi.org/10.1073/pnas.1603992113

44. Whitehead A, and Crawford DL. (2005). Variation in tissue-specific gene expression among natural populations. Genome biology, 6(2), 1–14. https://doi.org/10.1186/gb-2005-6-2-r13

45. Yip AM, and Horvath S. (2007). Gene network interconnectedness and the generalized topological overlap measure. BMC bioinformatics, 8(1), 1–14. https://doi.org/10.1186/1471-2105-8-22

46. Yoshihama M, Uechi T, Asakawa S, Kawasaki K, Kato S, Higa S, … and Kenmochi N. (2002). The human ribosomal protein genes: sequencing and comparative analysis of 73 genes. Genome research, 12(3), 379–390. http://www.genome.org/cgi/doi/10.1101/gr.214202

47. Yu K, Chen B, Aran D, Charalel J, Yau C, Wolf DM, et al. (2019). Comprehensive transcriptomic analysis of cell lines as models of primary tumors across 22 tumor types. Nature communications, 10(1), 1–11. https://doi.org/10.1038/s41467-019-11415-2

48. Zhang W, and Liu HT. (2002). MAPK signal pathways in the regulation of cell proliferation in mammalian cells. Cell research, 12(1), 9–18. https://doi.org/10.1038/sj.cr.7290105

49. Zoubarev A, Hamer KM, Keshav KD, McCarthy EL, Santos JRC, Van Rossum T, et al. (2012). Gemma: a resource for the reuse, sharing and meta-analysis of expression profiling data. Bioinformatics, 28(17), 2272–2273. https://doi.org/10.1093/bioinformatics/bts430

