## Supplementary Files for "Elucidating gene expression patterns across multiple biological contexts through a large-scale investigation of transcriptomic datasets"

### Supplementary File

#### Supplementary Text

**Supplementary Text 1. Applications of controllability analysis to biological networks**
A study by Vinayagam *et al.* (2016) applied this controllability analysis to a human PPI network, finding that 21% of the nodes in their human interactome were indispensable. Their results found that often, the indispensable nodes of a biological network are those targeted by viruses, disease-causing mutations, or are known drug targets. The findings indicate that network control can be a fundamental factor to the healthy vs disease state in humans. This was further investigated using data from over 1,500 cancer patients in which frequently altered genes in the disease state of nine different cancers were often indispensable nodes, of which 80% of those genes were not formerly known to be cancer-associated. This shows that controllability analysis in network biology can be a critical tool in discovering novel disease genes and finding new drug targets. In order to investigate interesting proteins such as potential novel disease genes and drug targets in the context-specific co-expression networks, we leveraged the results of the controllability analysis on the human PPI interactome as the co-expression networks are undirected.

**Supplementary Text 2. Cutoffs for most and least common nodes across contexts.**
For the most common proteins in each context, we used cut-offs from ≥ 46-43 out of 46 to create four groups ranging from 10 to 400 of the most common proteins in the tissue context. In the cell type context, we used cut-offs from ≥ 30-26 out of 30 to create four groups ranging from 150 to 1,800 of the most common proteins. In the cell line context, we used cut-offs from ≥ 22-19 out of 22 to create four groups ranging from 100 to 2,200 of the most common proteins. For the least common proteins in each context, we used cut-offs from ≤ 15-21 out of 46 to create four groups ranging from 100 to 400 of the least common proteins in the tissue context. In the cell type context, we used cut-offs from ≤ 9-13 out of 30 to create four groups ranging from 100 to 400 of the least common proteins. In the cell line context, we used cut-offs from ≤ 2-5 out of 22 to create four groups ranging from 75 to 400 of the least common proteins. The resulting proteins found in the overlap of each of these groups with the interactome can be seen in **Supplementary Table 8**.

**Supplementary Text 3. Most common edges for each context.**
In the tissue networks, we found 25 edges in more than 78% of the tissues (36/46) and 1,481 in more than 50% of the tissues (23/46). In the cell type networks, there were 25 edges in at least 70% of the cell types (21/30) and 962 in 50% (15/30) or more. In the cell line networks, we found 28 edges in more than 60% of the cell lines (13/22) and 354 in more than 50% (11/22).

### Supplementary Figures

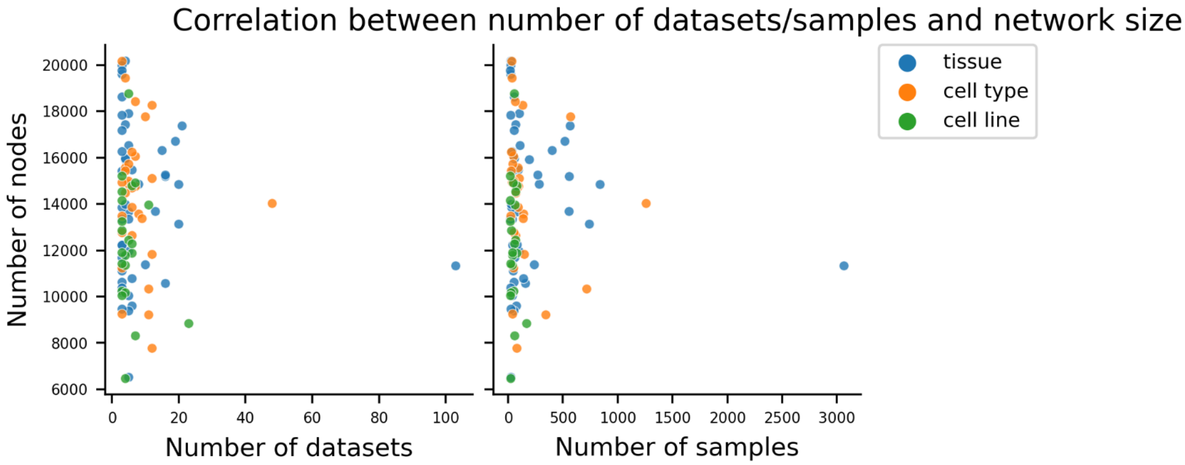


**Supplementary Figure 1. Correlation between number of nodes to number of datasets (left) and number of samples (right) for all networks.** The total number of datasets ranged from 3 to 103 in the tissue context, 3 to 48 in the cell type context, 3 to 23 in the cell line context. While the total number of samples was between 20 and 3,064 for the tissue context, 24 and 1,259 in the cell type context, 20 and 169 in the cell line context. The vast majority of co-expression networks were generated from 1 to 10 datasets and contained between 9 to 500 samples. We found that the resulting network size for each network varied within a wide range (i.e., between ~6,000 and 20,000) and no discernible pattern was observed.

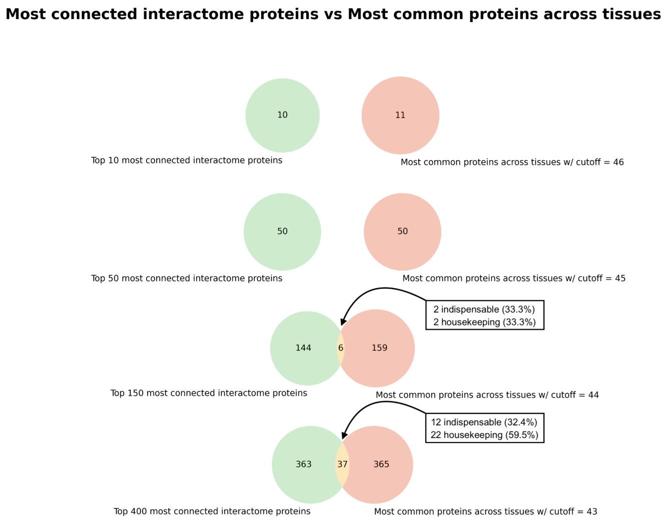


**Supplementary Figure 2. Overlap between the most common proteins of the tissue networks and the most connected proteins of the interactome.** Of the proteins present in the interactome and at least one tissue network, those of which are the most highly connected in the interactome (green) and most common in the majority of tissue co-expression networks (red), are compared at several scales (i.e., 10-400 proteins). In the overlap (yellow), the number of proteins that are indispensable in the interactome or housekeeping genes are stated. We find that, despite an increasing number of proteins analyzed, little overlap is observed.

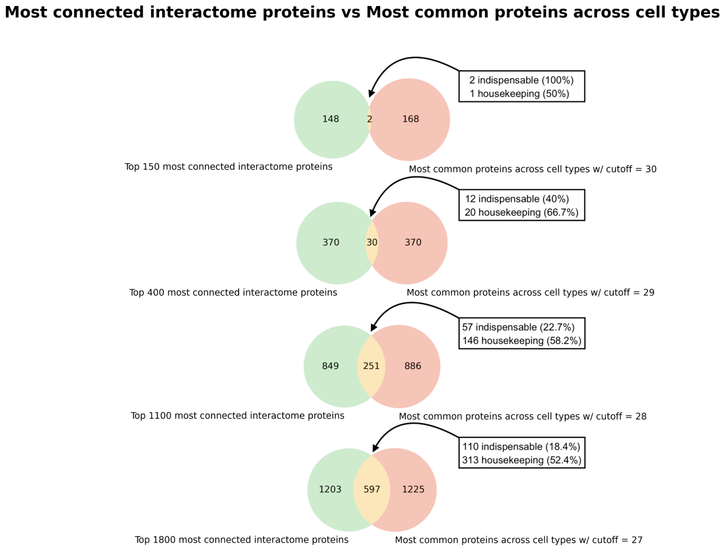


**Supplementary Figure 3. Overlap between the most common proteins of the cell type networks and the most connected proteins of the interactome.** Of the proteins present in the interactome and at least one cell type network, those of which are the most highly connected in the interactome (green) and most common in the majority of cell type co-expression networks (red), are compared at several scales (i.e., 150-1,800 proteins). In the overlap (yellow), the number of proteins that are indispensable in the interactome or housekeeping genes are stated. We find that, increasing the number of proteins analyzed, a significantly increasing overlap is observed.

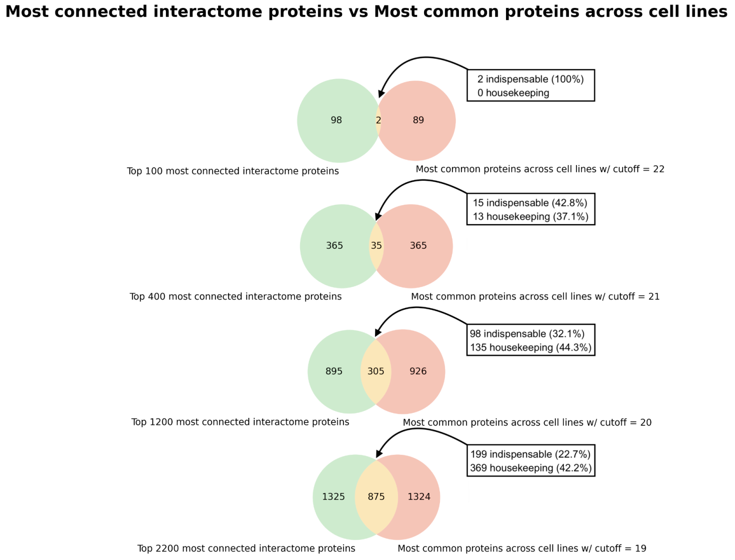


**Supplementary Figure 4. Overlap between the most common proteins of the cell line networks and the most connected proteins of the interactome.** Of the proteins present in the interactome and at least one cell line network, those of which are the most highly connected in the interactome (green) and most common in the majority of cell line co-expression networks (red), are compared at several scales (i.e., 100-2,200 proteins). In the overlap (yellow), the number of proteins that are indispensable in the interactome or housekeeping genes are stated. We find that, increasing the number of proteins analyzed, a significantly increasing overlap is observed.

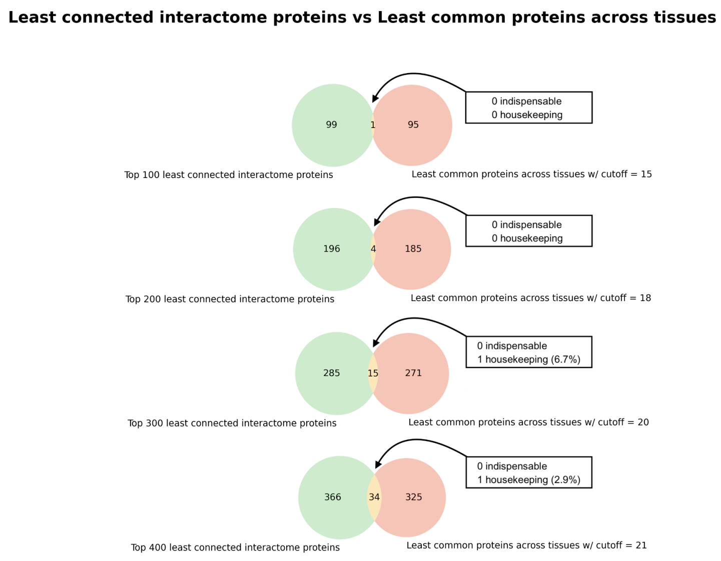


**Supplementary Figure 5. Overlap between the least common proteins of the tissue networks and the least connected proteins of the interactome.** Of the proteins present in the interactome and at least one tissue network, those of which are the least connected in the interactome (green) and least common in the majority of tissue co-expression networks (red), are compared at several scales (i.e., 100-400 proteins). In the overlap (yellow), the number of proteins that are indispensable in the interactome or housekeeping genes are stated. We find that, despite an increasing number of proteins analyzed, little overlap is observed.

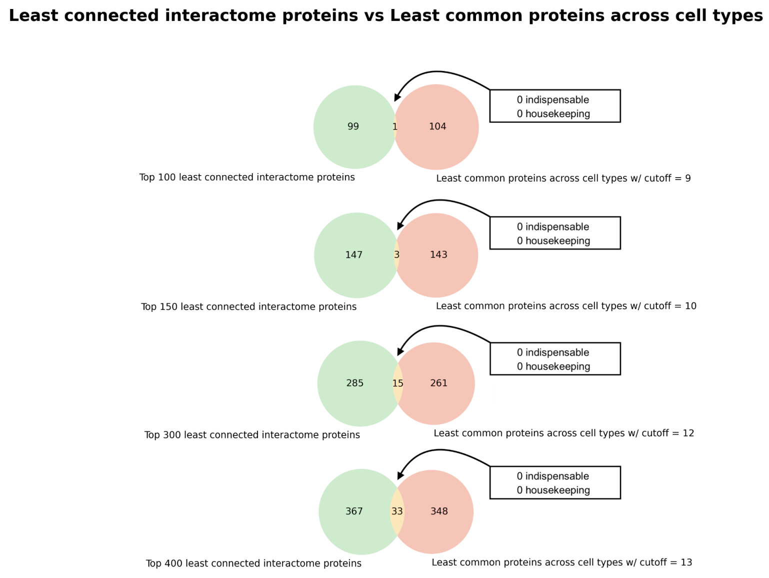


**Supplementary Figure 6. Overlap between the least common proteins of the cell type networks and the least connected proteins of the interactome.** Of the proteins present in the interactome and at least one cell type network, those of which are the least connected in the interactome (green) and least common in the majority of cell type co-expression networks (red), are compared at several scales (i.e., 100-400 proteins). In the overlap (yellow), the number of proteins that are indispensable in the interactome or housekeeping genes are stated. We find that, despite an increasing number of proteins analyzed, little overlap is observed.

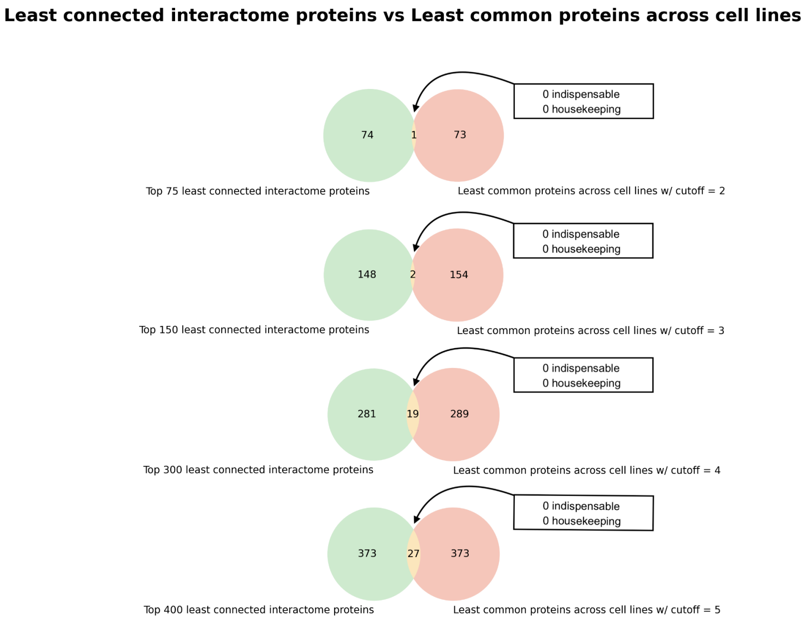


**Supplementary Figure 7. Overlap between the least common proteins of the cell line networks and the least connected proteins of the interactome.** Of the proteins present in the interactome and at least one cell type network, those of which are the least connected in the interactome (green) and least common in the majority of cell type co-expression networks (red), are compared at several scales (i.e., 75-400 proteins). In the overlap (yellow), the number of proteins that are indispensable in the interactome or housekeeping genes are stated. We find that, despite an increasing number of proteins analyzed, little overlap is observed.

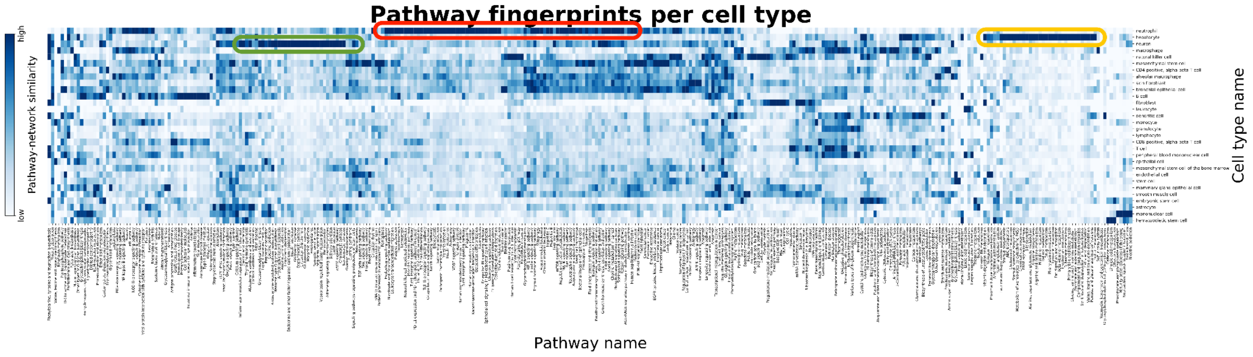


**Supplementary Figure 8. Similarity between cell type-specific co-expression networks and KEGG pathways.** The similarity between a particular pathway and a co-expression network is defined as the percentage of pairwise combinations of proteins of a given KEGG pathway that can be found in a co-expression network as edges. Light blue corresponds to a lower similarity, while dark blue corresponds to a high similarity. A high quality version of this figure is available at https://github.com/ContNeXt/scripts/blob/main/figures/supp_figure8_highquality.pdf and can also be visualized in the web application.

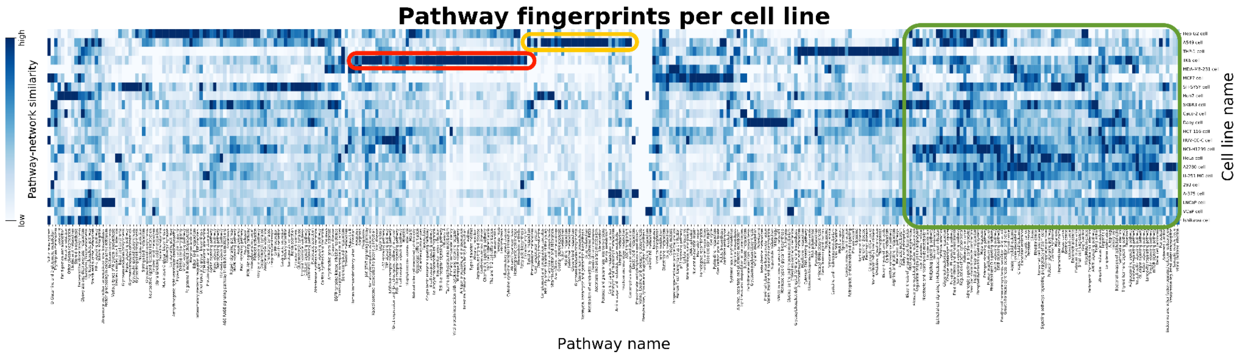


**Supplementary Figure 9. Similarity between cell line-specific co-expression networks and KEGG pathways.** The similarity between a particular pathway and a co-expression network is defined as the percentage of pairwise combinations of proteins of a given KEGG pathway that can be found in a co-expression network as edges. Light blue corresponds to a lower similarity, while dark blue corresponds to a high similarity. A high quality version of this figure is available at https://github.com/ContNeXt/scripts/blob/main/figures/supp_figure9_highquality.pdf and can also be visualized in the web application.
